# Altering sensory cues for spatial navigation does not impose a dual-task effect on gait and balance

**DOI:** 10.64898/2026.03.16.712118

**Authors:** Sam Beech, Maggie K. McCracken, Christina Geisler, Leland E. Dibble, Colby R. Hansen, Sarah H. Creem-Regehr, Peter C. Fino

## Abstract

Walking is an attentionally demanding process that draws from a limited pool of attentional resources. Dual-task assessments, where individuals perform a cognitive task while walking, often reveal changes in gait and balance due to competing attentional demands. As cognitive task difficulty increases, the attentional resources necessary to complete the task also increase, leading to greater interference with gait and balance. However, these interactions are typically examined using contrived lab-based tasks, leaving it unclear how the cognitive processes engaged during real-world movement impact walking. In the present study, we investigated whether increasing the attentional demand of spatial navigation, a cognitive process intrinsically linked to movement, interferes with gait and balance. Healthy adults completed an ambulatory virtual reality homing task in which they walked through a virtual environment and navigated to previously visited locations while wearing ankle and lumbar trackers. We increased the attentional demand of navigation by removing sensory cues during this homing phase: full cues, visual cues only, or self-motion cues only. Navigation performance declined as sensory cues were removed, but we observed no corresponding changes in their spatiotemporal gait and balance metrics. These results show that, in healthy adults, increasing the attentional demand of spatial navigation does not interfere with gait and balance during real-world movement. This finding suggests that locomotor control may be robust to navigation-related cognitive demands. Further research is needed to determine why navigation did not interfere with mobility and to clarify the relationship between these two interconnected processes.

## INTRODUCTION

Walking is a complex motor control process that requires estimating the body’s center of mass (Tesio & Rota, 2019), selecting appropriate foot placements to guide movement direction while maintaining balance (Bruijn & Van Dieën, 2018; Dingwell & Cusumano, 2019; Redfern & Schumann, 1994), and continuously repeating these processes using prior and incoming sensory feedback (Hausdorff, 2007; Rossignol et al., 2006). Yet walking rarely occurs in isolation; people typically engage in additional cognitive tasks while moving. The impact of concurrent cognitive engagement on locomotor control is commonly examined using dual-task paradigms, in which participants perform a cognitive task during movement. In these contexts, cognitive and motor processes draw from a shared, limited pool of attentional resources. When the combined attentional demands of the two tasks exceed capacity, neither receives sufficient resources, leading to measurable changes in gait and balance such as reduced stride speed (Al-Yahya et al., 2011; Howell et al., 2019; Kelly et al., 2010; Yogev-Seligmann et al., 2008; Woollacott & Shumway-Cook, 2002). Dual-task assessments are therefore widely used as clinical tools to identify locomotor deficits in older adults (Gobbo et al., 2014) and in populations with neurological conditions such as mild traumatic brain injury (mTBI) (Fino et al., 2016, 2018), stroke (He et al., 2018), and Alzheimer’s disease (Fritz et al., 2015). Although these assessments are valued for their ecological validity, they typically rely on contrived cognitive tasks – such as mental arithmetic or the Stroop task (Fino et al., 2016; Hiyamizu et al., 2012; McPhee et al., 2022; Rochester et al., 2014; Tsai et al., 2009; Ward et al., 2022; Worden et al., 2016) – that are uncommon during everyday mobility and unrelated to the goals of natural movement. More naturalistic tasks, such as extemporaneous speech, have recently been employed to assess deficits in individuals with mTBI (Yang et al., 2025; Poretti et al., 2025), but even this fails to fully capture the cognitive processes engaged in everyday locomotion. As a result, how real-world cognitive demands interfere with gait and balance during daily movement remains poorly understood.

Critically, walking in daily life rarely occurs without purpose; people typically move with the goal of reaching a desired location. Therefore, one cognitive process that is inherently linked to locomotion is spatial navigation, and the specific component of navigation, *spatial updating* – the process of maintaining an allocentric representation of one’s position within the environment during movement. Effective spatial updating requires attending to spatially relevant body-based and environmental information (Janzen & Van Turennout, 2004; Ruddle et al., 2013), processing this sensory input (Wolbers & Hegarty, 2010), mapping this information onto stored spatial representations (Epstein et al., 2007), and choosing the most efficient route to a destination (Patai & Spiers, 2021; Varshney et al., 2024). Broadly, information about one’s position within the environment is derived from two sets of cues. First, body-based self-motion cues, such as proprioceptive feedback and vestibular signals, provide cumulative translational and rotational information to support estimations of the distance and direction of travel (Cullen, 2019; Lappe et al., 1999; Loomis et al., 1993). Second, external visual landmarks provide immediate information about position and orientation relative to the surrounding environment (Bates & Wolbers, 2014; Chan et al., 2012; Nardini et al., 2008; Newman et al., 2023; Shayman et al., 2024). These redundant sensory cues are combined into a single coherent estimate of position and orientation relative to the world around us (Chen et al., 2017; Cheng et al., 2007; Nardini et al., 2008; Sjolund et al., 2018). Despite the close interdependence of locomotion and navigation, it remains unclear how these processes compete for limited attentional resources during daily movement.

Spatial updating is an active, attentionally demanding process. Greater cortical activation during navigation demonstrates its reliance on attentional resources (Do et al., 2020), which is also reflected behaviorally, as participants explicitly instructed to attend to and remember spatial locations during locomotion show enhanced recall later (Van Asselen et al., 2006). Consistent with this, introducing an unrelated cognitive task diverts attention away from navigation and impairs performance. Early work showed that learning an environmental layout and identifying key spatial reference points is significantly disrupted by concurrent reverse counting (Lindberg & Gärling, 1982; Smyth & Kennedy, 1982). Both navigational difficulty and cognitive-task difficulty further modulate the degree of interference. For example, performance on a simple single-leg path retracing task is comparably impaired by a low-demand verbalization task and a high-demand reverse-counting task, but when participants complete a more attentionally demanding three-leg triangle-completion task, the reverse-counting task produces significantly greater navigational decline (May & Klatzky, 2000). As attentional demands increase in either domain, interference grows because the total required resources for efficient performance increasingly exceed the attentional capacity.

Dual-task interference also increases when the attentional demands of navigation are increased by reducing the reliability of sensory cues that support spatial orientation. Klatzky et al. (2006) demonstrated this effect by comparing navigation performance and N-back accuracy when participants navigated using either spatial language (left/right/straight) or virtual sound (binaural cues signaling distance and direction in the yaw plane). Although single-task navigation performance did not differ between the two modalities, dual-task performance was superior in the virtual sound condition. This finding aligns with earlier work showing that binaural auditory cues yield more precise spatial information and are less attentionally demanding than spatial-language guidance (Loomis et al., 1998; Loomis et al., 2005). Similarly, degrading traditional sensory cues also increases interference, as performance on an auditory reaction-time task worsens when participants navigate with blurred vision compared to normal vision (Rand et al., 2015). Together, these findings show that decreasing the reliability of spatial information increases the attentional demands of navigation, thereby increasing interference with concurrent cognitive tasks. Degradation of the sensory cues that facilitate navigation is common in everyday mobility, such as changing compliance in the terrain, indistinguishable landmark configurations, or low-light conditions at night, but how these environmental changes impact gait and balance during movement remains unclear.

In dual-task walking paradigms, increasing the complexity of the concurrent cognitive task similarly exacerbates the interference with gait and balance. Young healthy adults exhibit greater whole-body angular momentum – a marker of reduced balance control (Neptune & Vistamehr, 2019) – when reverse-spelling long versus short words while walking (Small et al., 2021). Similarly, both young and older adults walk more slowly when completing a cognitively demanding clock-matching task (hearing a time and judging whether the hour and minute hands fall within the same left/right half of an observed clock face) compared to a simpler auditory Stroop task (Plummer-D’Amato et al., 2012). Reductions in gait speed have also been observed with increased difficulty in backward counting, list recall, reaction-time, or word-generation tasks (Goh et al., 2021). As cognitive-task complexity rises, the number of attentional resources required to perform that task also increases, resulting in greater interference with locomotor control.

Although navigation and balance in real-world settings have typically been examined as separate processes, there is growing recognition and evidence of their interaction and potential interference. This potential interaction has been leveraged in clinical research to model gait characteristics during navigation for classifying young versus older adults (Pawlaczyk et al., 2021) and for differentiating older adults with and without medial temporal lobe atrophy (Pawlaczyk et al., 2023). The first direct investigation of dual-task interference between navigation and locomotion was conducted by Rand et al. (2015), who manipulated the attentional demands of walking by allowing healthy young adults to navigate either with or without physical support from the experimenter (holding their arm). Landmark recall was significantly higher when participants walked with physical support, suggesting that the reduced locomotor demands freed attentional resources for navigation. Older adults (60+ years) also displayed superior landmark recall during navigation in a novel environment when walking with physical support from the experimenter (Barhorst-Cates et al., 2017). The reverse relationship has also been observed: increased navigational difficulty can impair gait. Camp et al. (2023) investigated the impact of increasing the difficulty of The Walking Corsi Test (WCT) on gait. The WCT involves a configuration of nine landmarks (0.45 × 0.45 m) arranged in a non-uniform pattern within the testing space (4.5 × 3.75 m). The participants were seated as they observed an experimenter walk in sequence to the landmarks. They then had to walk to these landmarks in standard or reverse order. As the WCT sequence number increased, participants exhibited greater changes in step time, stance time, stride speed, and cadence, indicating interference from navigational demands. However, the WCT primarily assesses visuospatial working memory and serial-order reproduction based solely on visual observation (Nori et al., 2015), limiting its ecological validity. Real-world navigation, by contrast, relies on the continuous updating of one’s position relative to an allocentric reference frame through both visual feedback and path integration – the use of proprioceptive, vestibular, and optic flow signals to update self-estimates of location during movement. The observation of disrupted locomotor control as the WCT became more challenging is consistent with previous dual-task balance paradigms. However, it remains unclear whether increasing the difficulty of navigation in an ecologically valid context, where one must continuously monitor and estimate their own position relative to the world during movement, interferes with gait and balance.

Therefore, the purpose of this study was to determine whether increasing the attentional demands of navigation through the removal of spatially informative sensory cues interferes with gait and balance during ambulatory navigation. To test this, we used an immersive virtual reality (VR) homing task that enabled precise control over environmental cues and continuous measurement of locomotor behavior. Participants navigated a virtual environment (VE) under three sensory-cue conditions (Full Cues, Vision Only, and Self-Motion Only) that systematically varied the availability of spatial information (Shayman et al., 2024). As participants walked, we collected a range of gait, balance, and navigation performance metrics. To assess whether increasing navigation difficulty interferes with locomotor control, we compared performance in the Full Cues condition with the two single-cue conditions. As reducing the reliability of the spatial cues available for navigation has previously been shown to increase attentional demand (Rand et al., 2015), we hypothesized that the two single-cue conditions would elicit greater interference with gait and balance, producing measurable changes in step length, step width, and stride speed, relative to the Full Cues condition.

## MATERIALS and METHODS

### Participants

The present experiment utilized a subset of participants from two ongoing studies that employed the same VR homing task to investigate cue combination during sensory integration for navigation in populations with compromised sensory processing. The first study focuses on individuals with mild traumatic brain injury, while the second examines individuals experiencing varying degrees of simulated peripheral visual field loss. For the current analysis, we included 24 healthy control participants from study one and 8 healthy control participants from study two. Of the 32 total participants, 22 identified as female (Mean age = 22.8; SD = 5.9) and 10 identified as male (Mean age = 24, SD = 4.6). Participants were required to be aged between 18 and 50 years old, have normal or corrected-to-normal vision, and be able to walk independently. Exclusion criteria included a concussion within the past year or a history of three or more lifetime concussions; any current or past neurological, musculoskeletal, or vestibular conditions affecting balance; a history of heart conditions or seizures; current pregnancy; or meeting criteria for moderate-to-severe substance use disorder within the past month as defined by the DSM-5. Participants whose behavior posed a safety concern or compromised data integrity were also excluded. All participants provided written informed consent prior to participation, and all procedures were approved by the Institutional Review Board at the University of Utah (IRB_00166649, IRB_00156396).

### Procedures and design

Participants were fitted with an HTC Vive Pro Eye head-mounted display (HMD) and Vive Trackers 3.0 were placed over the ankles (lateral malleoli) and the lumbar (L5/S1) junction. The headset and trackers provided three-dimensional position and orientation data aligned with the VE dimensions. Brown noise was played through earphones to minimize auditory spatial cues. The experimenter informed the participant that they would be accompanied throughout the experiment and stopped if they approached a boundary, allowing them to walk naturally without concern for real-world collisions.

The VE was rendered on the HMD prior to setup so that participants were immediately immersed in the virtual testing space (Figure 1). The participants were instructed to walk to the starting point (gold star - Figure 1) and then turn to face the markers and landmarks. In the tracing phase, the participants were instructed to walk to the ground markers in a set sequence: black-red-blue. Upon reaching the blue marker, the HMD blacked out for ten seconds, and the ground markers were removed.

**Figure 1:**
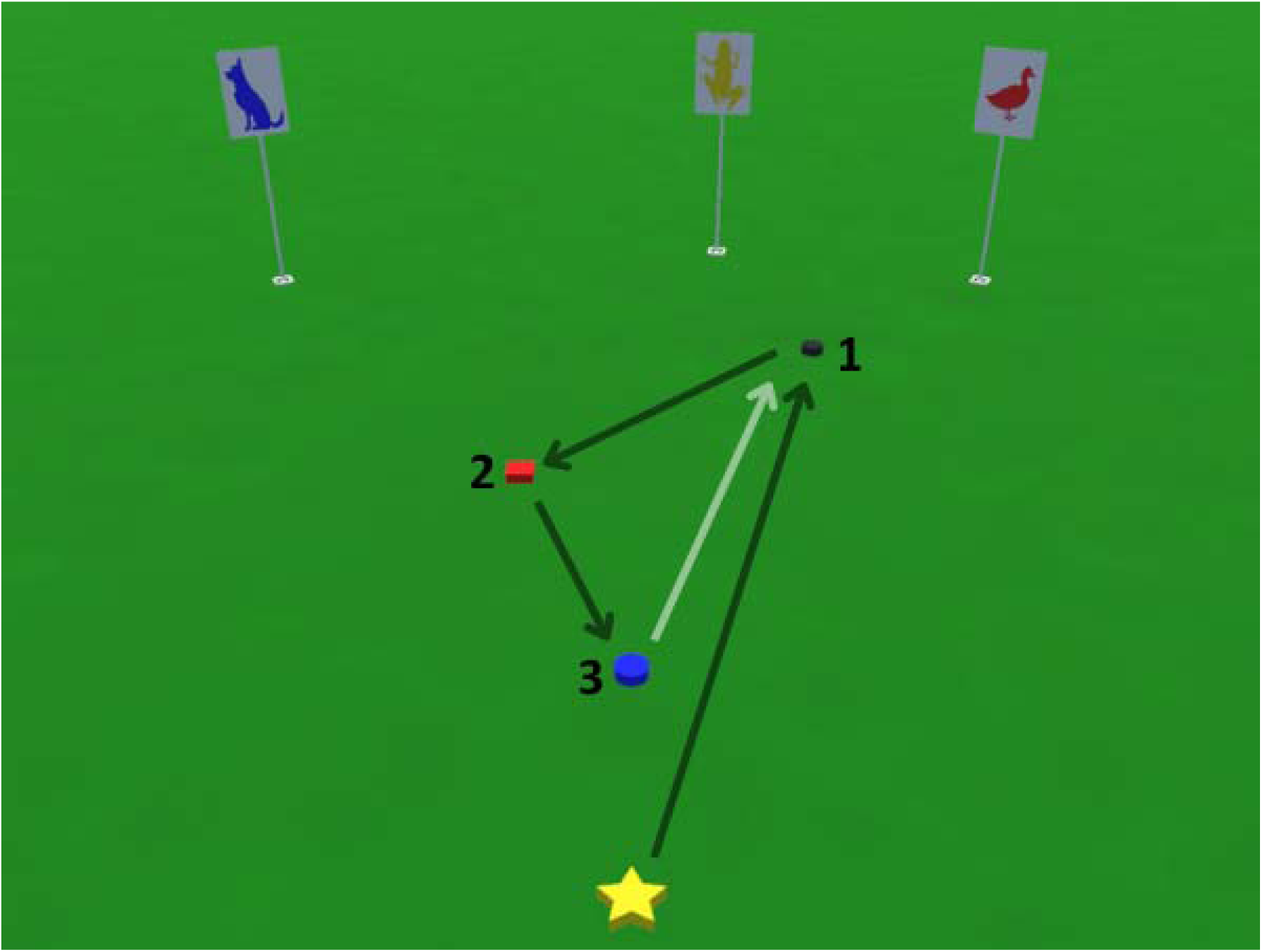
Illustration of an example trial. Participants begin at the gold star and walk to the three ground markers in sequence with the landmarks displayed in the background. At the blue marker, the ground markers are removed, and the participant must then walk to their estimates position of the black marker,

Participants were instructed to remain still during this blackout period. When the display re-illuminated, the participants were asked to walk to their estimated position of the black marker. The homing phase was performed in one of four conditions: **Full cues** – no manipulation to sensory feedback during the blackout; **Self-Motion Only** – visual landmarks were removed during the blackout, leaving only vestibular and proprioceptive cues to estimate spatial location; **Vision Only** – visual landmarks remained visible, but participants were rotated in a chair at a quick but gentle speed, with rotations occurring equally in both directions for 10 seconds during the blackout to disrupt vestibular and proprioceptive cues and disorient their ‘sense of direction’. Participants were not aware that they were spun equally in both directions and returned to their initial orientation, as the spinning comprised of smaller clockwise and counterclockwise rotational components. This approach has been validated in previous research (Bates & Wolbers, 2014; Chen et al., 2017; Shayman et al., 2024); and **Conflicting Cues** – landmarks were shifted 15° to the left relative to the starting position during the blackout to create spatial conflict between the self-motion-defined (unshifted) position and the landmark-defined (shifted) position (Figure 2). The role of the Conflicting Cues condition was to identify sensory weighting for navigation. Therefore, as this was not an aim of the present study, and as the impact of Conflicting Cues on attentional demand and locomotor behavior is less defined, this condition was not included in our current analysis. For each trial, the participants confirmed their estimated position of the black marker by standing still. This position was then saved, and the headset was blacked out again. The experimenter then walked the participant back to the starting position and the headset re-illuminated. At the start of each trial, a verbal reminder of the black-red-blue sequence was provided.

**Figure 2:**
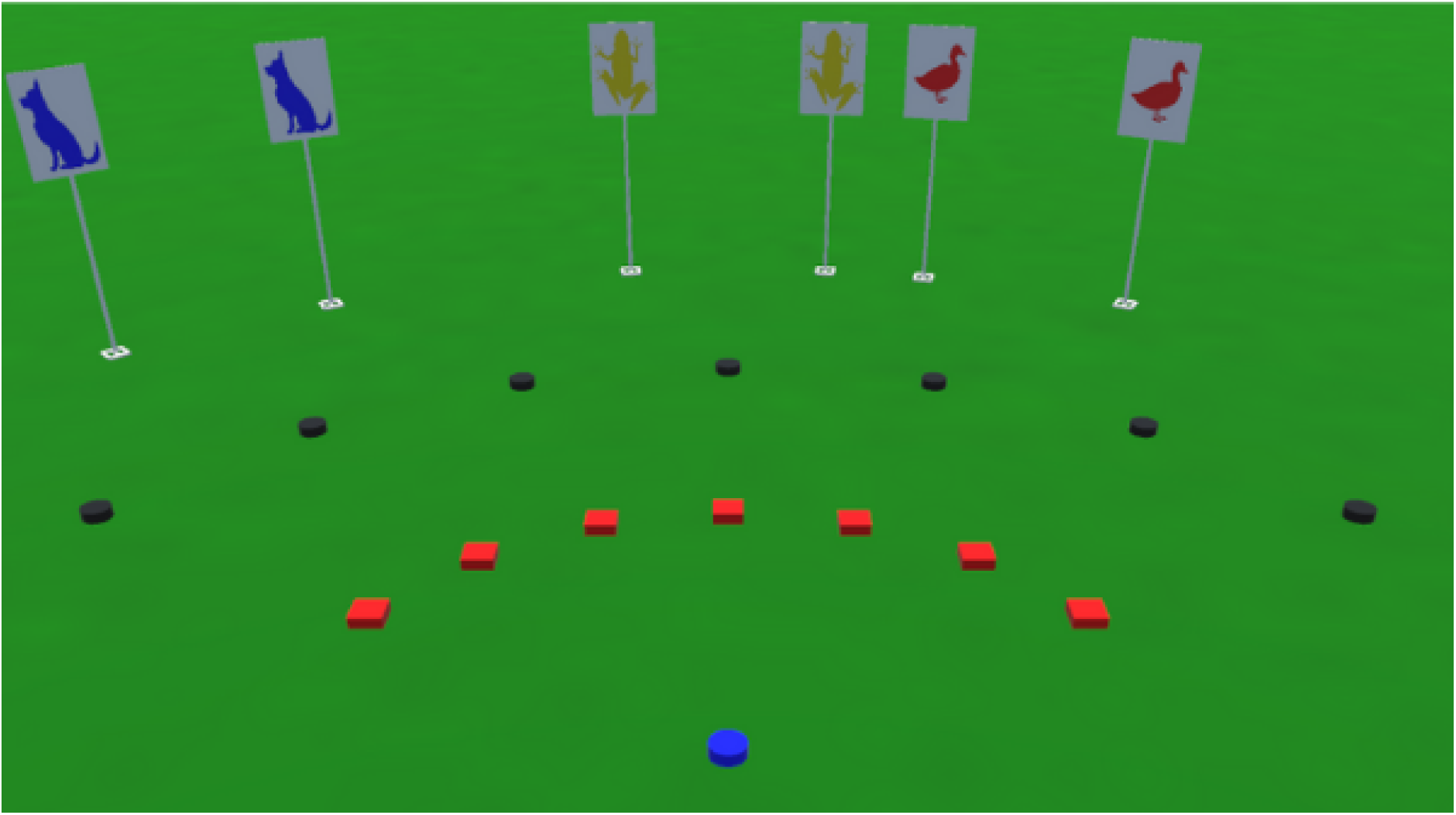
All potential configurations. The right-sided landmark represents the typical position throughout each tracing phase, and when presented in the Full Cues and Vision Only homing phases (as shown in Figure 1). The left-sided landmarks represents the shifted position after the blackout during the Conflict condition. The ground markers show all potential configutations for black and red markers (i.e., waypoints 1 and 2 from Figure 1), but the blue position (waypoint 3) remained unchanged.

The positions of the black and red ground markers changed between trials, but the blue marker was always located in the same location, just in front of the starting position (Figure 2). The positions of the landmarks during the tracing phase did not change throughout the experiment. The 24 participants from study one completed 9 trials in each condition while the 8 participants from study two completed 8 trials in each condition. However, the first trial in each condition was discarded as a practice trial, leaving 8 and 7 trials for analysis, respectively. Trials were also discarded if the experimenter had to stop the participant from walking into a real-world object, the experimenter saved the homing position too early, or the participant ignored instructions.

### Measured outcomes and data preprocessing

The orientation and position of the headset, lumbar, and ankle trackers were recorded at 90 Hz throughout the homing task. We collected various spatiotemporal gait metrics throughout the homing phase, such as: step width, calculated as the mediolateral component of the vector between the left and right ankle positions at each heel strike projected onto the mediolateral axis defined by the participant’s lumbar yaw orientation; step length, calculated as the anteroposterior component of the same vector projected onto the anteroposterior axis defined by the lumbar orientation; and stride speed, calculated as the sum of two consecutive step lengths divided by the time elapsed between heel strikes of the same limb. Navigation performance was quantified as the error, in meters, between the estimated and actual position of the black marker in the homing phase.

### Statistical Analysis

A set of linear mixed-effects models (LMMs) were used to investigate differences in the average step width and step length (and their variability), stride speed, and navigation accuracy across three homing conditions (Full Cues, Self-Motion Only, and Vision Only). Each model included a Satterthwaite approximation and F-tests for fixed effects, with homing condition specified as a categorical fixed effect and participant number included as a random-effects factor with random intercepts and slopes to account for variability across participants. All analyses were conducted in JASP (Version 0.19.1; University of Amsterdam, 2024) using a significance level of 0.05.

As frequentist analyses cannot provide evidence for the absence of an effect, Bayes factors (BF₀₁) were computed using the ‘*BayesFactor’* package in R V4.5.2 when analyzing the main effect of the homing condition. BF₀₁ represents an odds ratio quantifying how much more likely the observed data are under the null hypothesis (no Condition effect) than under the alternative hypothesis (Condition effect). Values greater than 3 indicate strong evidence in favor of the null hypothesis, whereas values less than 0.3 indicate strong evidence in favor of the alternative hypothesis (Kass & Raftery, 1995).

## RESULTS

The average spatial error between the estimated and actual position of the black marker was larger in the single-cue conditions (Figure 3). The model reported a significant main effect of condition, (F(2, 40.74) = 15.651, p < .001, BF₀₁ = 2.90 × 10⁻), and follow-up Holm-Bonferroni corrected contrast tests report that navigation error in the Full Cues condition was significantly smaller than the Self-Motion Only condition (β = 0.321, t(35.29) = 5.452, p < .001) and the Vision Only condition (β = 0.153, t(56.55) = 3.084, p = .006). Additionally, navigation error in the Vision Only condition was significantly smaller than the Self-Motion Only condition (β = 0.168, t(31.48) = 2.701, p = .011).

**Figure 3:**
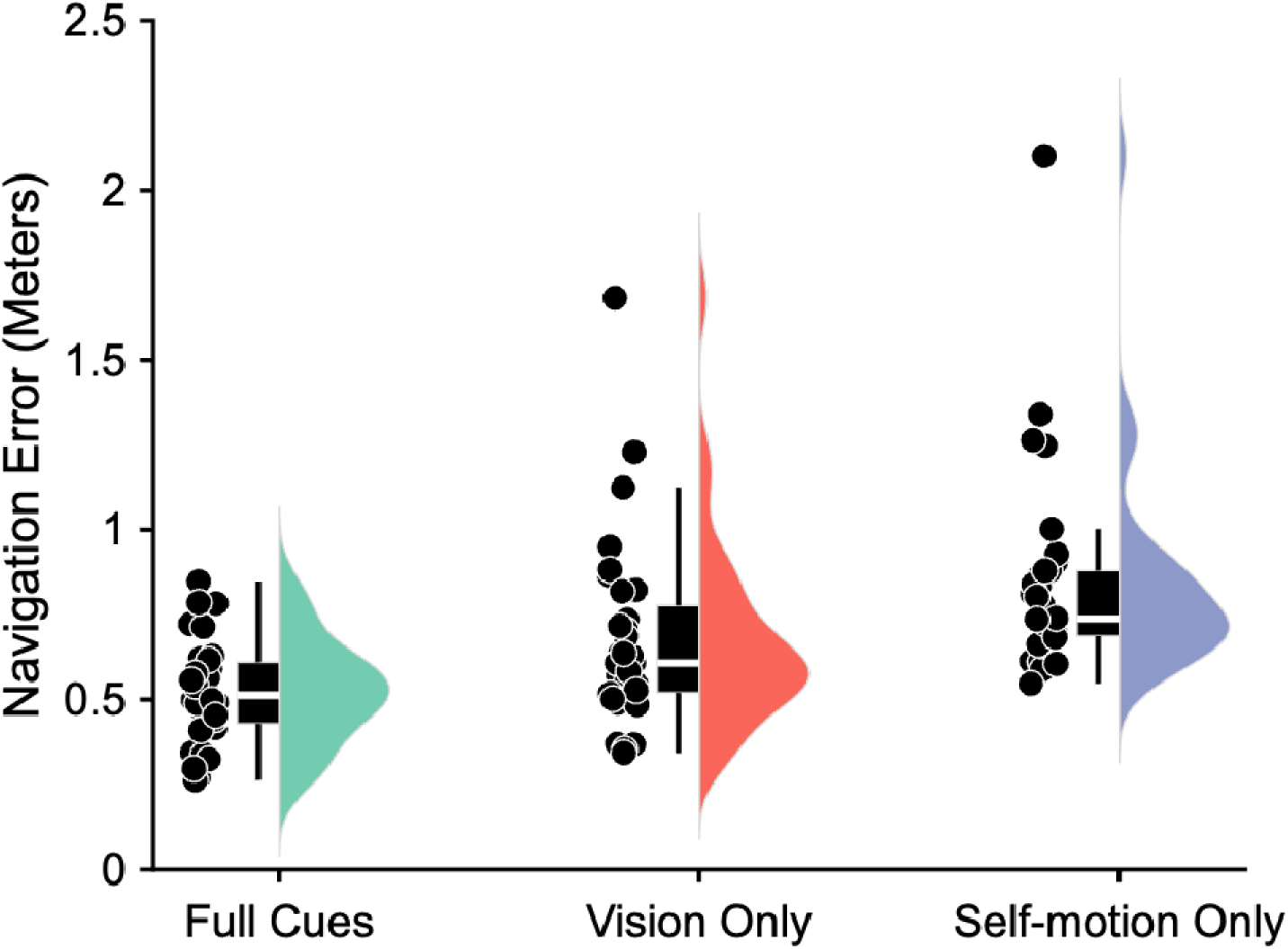
The average error between the estimated and actual black marker position for each homing condition. The scattered points represent individual participant averages. The violin plots illustrate the distribution, with box plots showing the median (white line) and quartile ranges (black box).

Although step length did not significantly change from the Full Cue to single-cue conditions, a significant main effect of condition was observed (F(2, 52.61) = 4.812, p = .012, BF₀₁ = 1.01). Holm-Bonferroni corrected contrast tests reported no difference from the Full Cues to the Self-Motion Only (β = 0.004, t(33.39) = 0.633, p = .531) or Vision Only condition (β = 0.012, t(38.21) = 2.12, p = .081), but step length in the Self-Motion Only condition was longer than the Vision Only condition (β = 0.016, t(192.36) = 2.949, p = .011). We also observed no effect of condition on step length variability (F(2, 53.03) = 0.009, p =. 991, BF₀₁ = 116.75).

**Figure 4:**
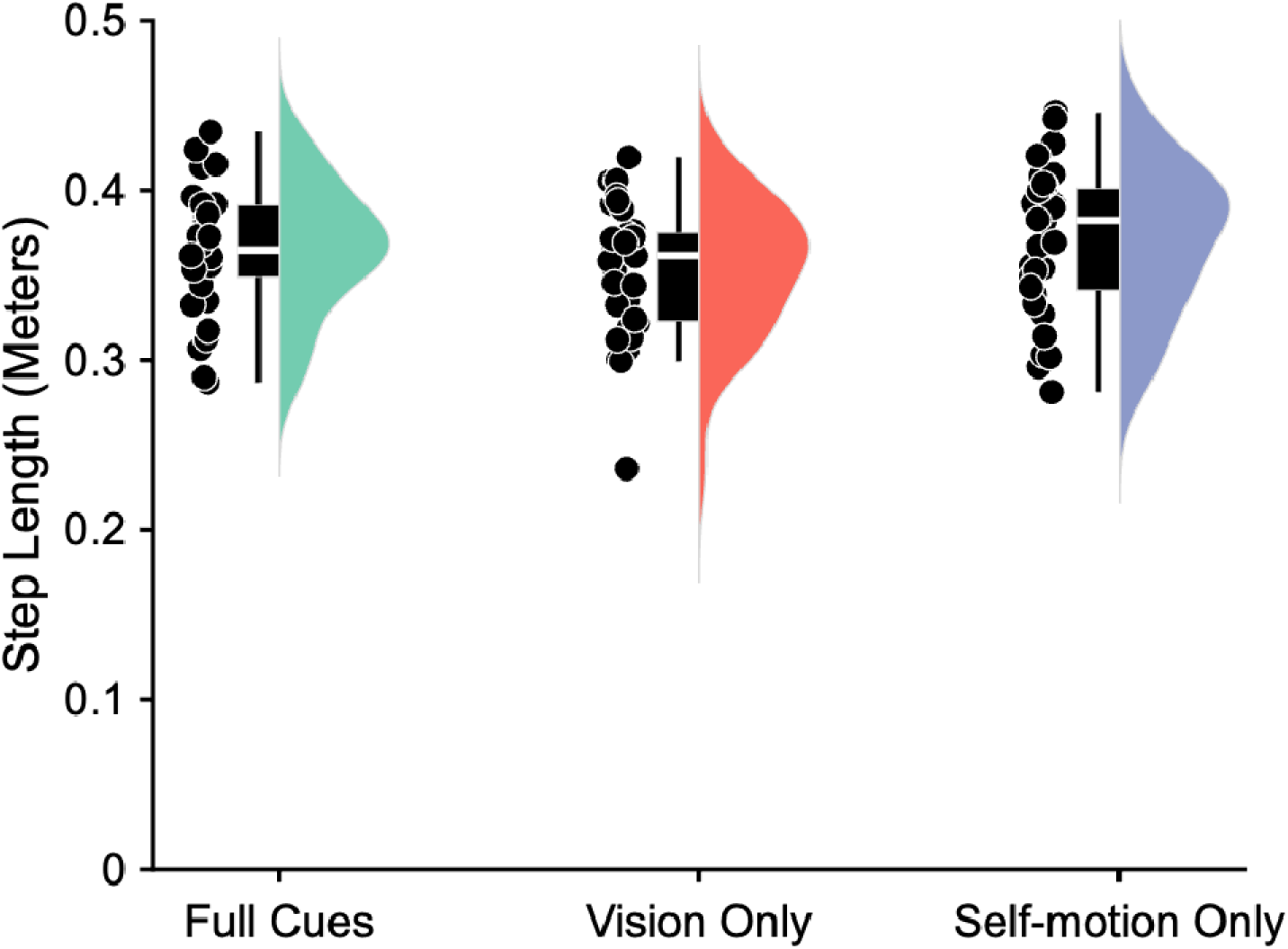
The average step length for each homing condition. The scattered points represent individual participant averages. The violin plots illustrate the distribution, with box plots showing the median (white line) and quartile ranges (black box).

Manipulation of the sensory cues during the homing phase had no effect on step width. There was no significant effect of condition for the average step width (F(2, 53.33) = 0.102, p =. 903, BF₀₁ = 107.10) or step width variability (F(2, 37.79) = 0.945, p =. 398, BF₀₁ = 43.65).

**Figure 5:**
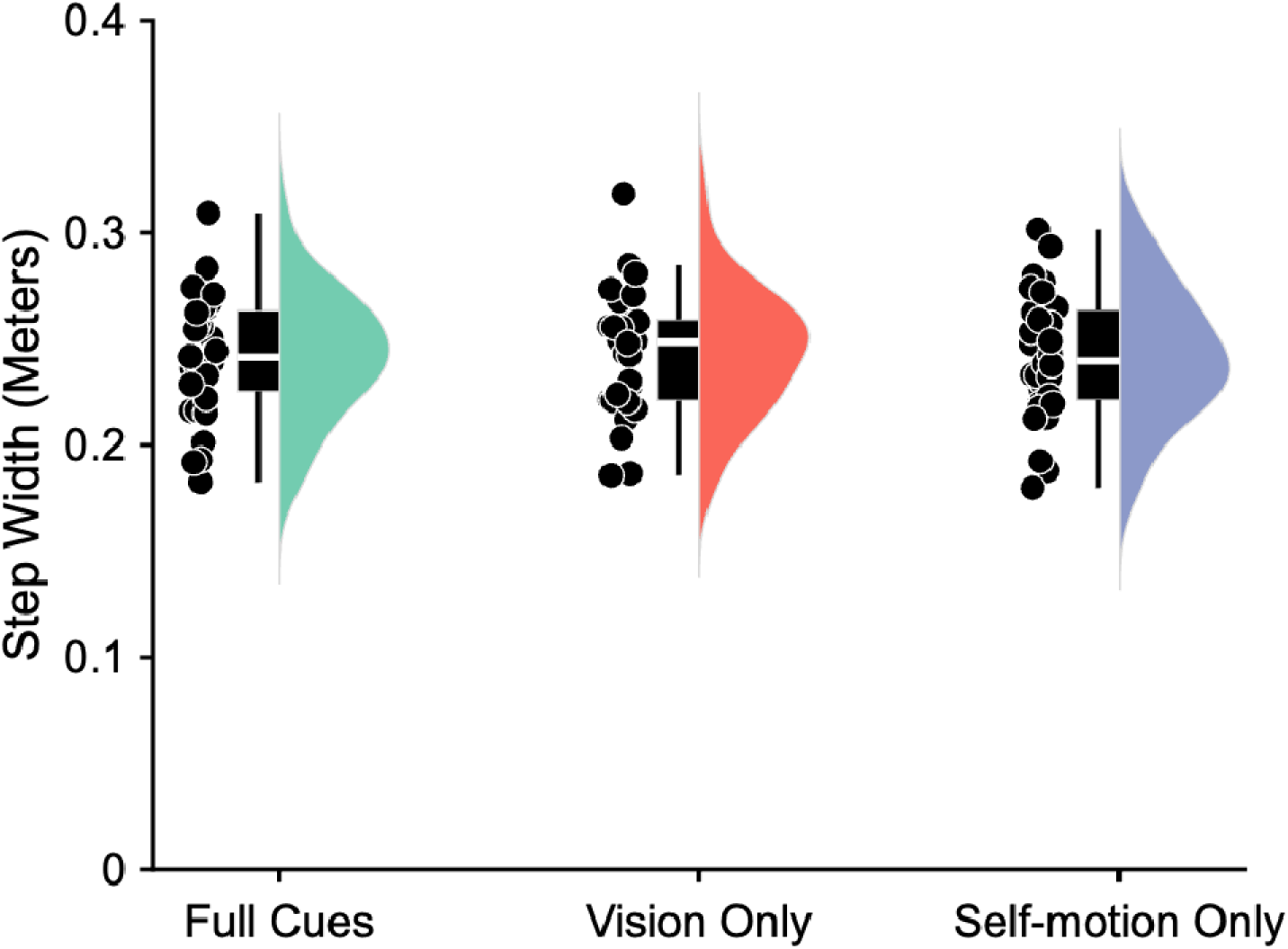
The average step width for each homing condition. The scattered points represent individual participant averages. The violin plots illustrate the distribution, with box plots showing the median (white line) and quartile ranges (black box).

Manipulation of the sensory cues during homing had no impact on stride speed. We observed no effect condition for the average stride speed (F(2, 65.21) = 1.822, p = .170, BF₀₁ = 16.54).

**Figure 6:**
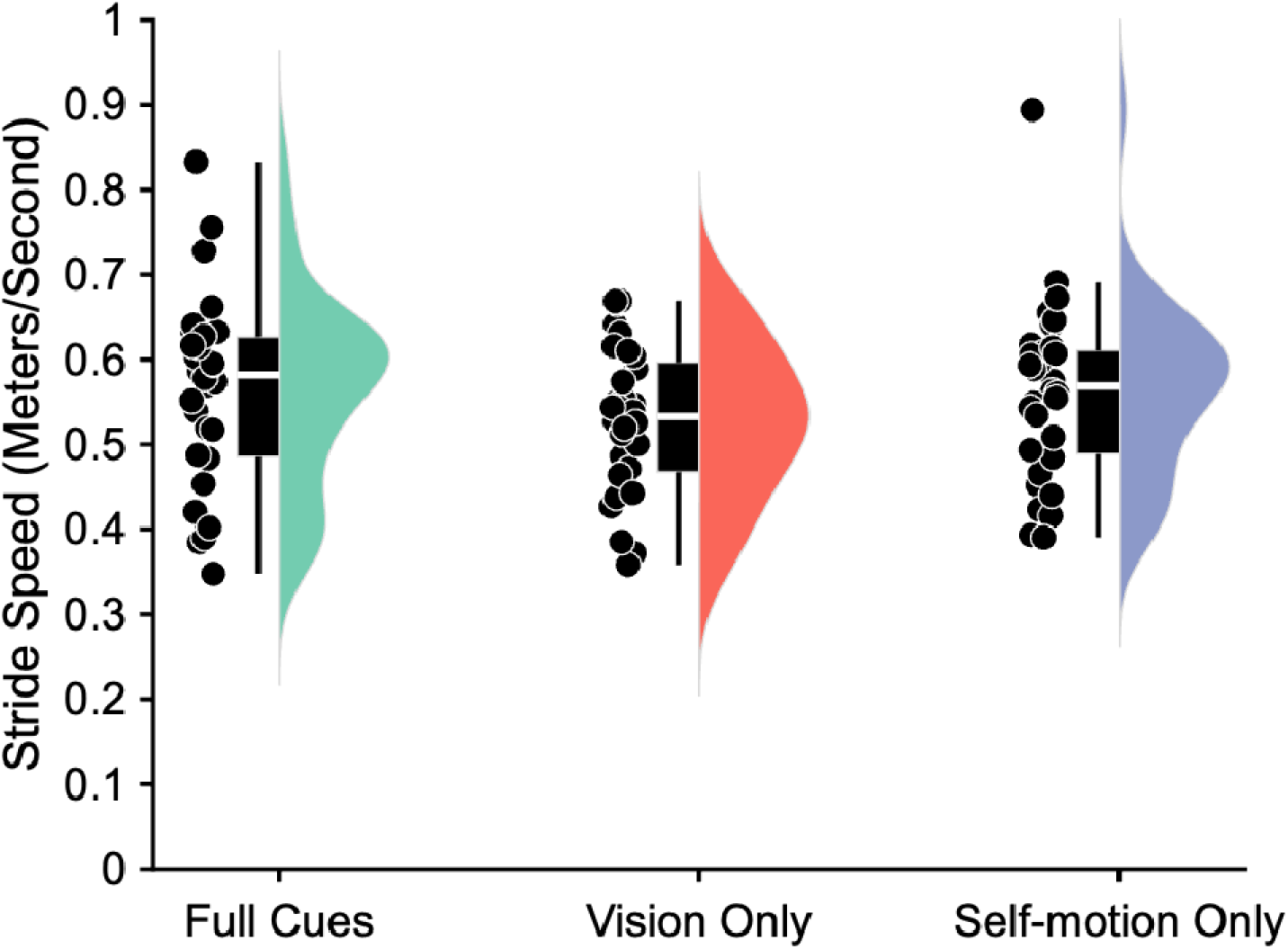
The average stride speed for each homing condition. The scattered points represent individual participant averages. The violin plots illustrate the distribution, with box plots showing the median (white line) and quartile ranges (black box).

## DISCUSSION

This study aimed to determine whether increasing the difficulty of navigation by removing spatially informative sensory cues from the VE would cause interference with gait and balance during an ambulatory VR homing task. We hypothesized that removing sensory cues would increase the attentional resources required for effective navigation and, in turn, lead to greater interference with gait and balance. However, we observed no significant differences in spatiotemporal gait metrics between the Full Cues and single-cue conditions, indicating that gait and balance were preserved despite the removal of these cues. In contrast, navigation error increased under both single-cue conditions. Therefore, increasing the difficulty of navigation impaired homing performance but did not disrupt locomotor performance.

One common explanation for preserved gait with declining performance in a concurrent cognitive task is the “posture-first strategy.” Originally proposed by Shumway-Cook et al. (1997), this ecological explanation suggests that individuals prioritize the primary task of maintaining postural stability over the less critical cognitive task. Specifically, Plummer & Eskes (2015) describes our pattern – declining cognitive performance coupled with unchanged balance – as “cognitive interference.” In dual-task walking paradigms, posture-first interpretations most often refer to smaller changes in gait relative to large changes in cognitive task performance (Hennah & Doumas, 2023; Pitts et al., 2023; Ruffieux et al., 2015). This finding is also observed following explicit prioritization, as even with instruction to focus on controlling gait, participants frequently show measurable changes such as increased walking velocity (Canning, 2005; Siu et al., 2008; Yogev-Seligmann et al., 2010) or reduced, but not eliminated, dual-task gait costs (Kelly et al., 2013; Verghese et al., 2007; Yogev-Seligmann et al., 2012). More challenging cognitive tasks further increase dual-task costs, with gait prioritization attenuating but not abolishing these effects (Maclean et al., 2017). In the present study, however, spatiotemporal gait metrics did not change from the Full Cue to single-cue conditions. This result could be verified as a posture-first strategy if single-task navigation performance remained unchanged across the different sensory cue conditions, but such a comparison was not possible because ambulatory navigation requires walking. Moreover, given that gait changes are commonly observed even when gait is explicitly prioritized, we do not believe the posture-first strategy wholly explains our results.

An alternative explanation for these results is that the sensory cue manipulations did not sufficiently increase the attentional demand of navigation to exceed attentional capacity. Prior work shows that navigating with less reliable cues can heighten attentional demand and produces larger dual-task costs (Klatzky et al., 2006; Rand et al., 2015), and that increasing the difficulty of a concurrent cognitive task typically generates greater interference with gait and balance (Goh et al., 2021; Plummer-D’Amato et al., 2012; Small et al., 2021). The increased navigation error observed in the single-cue conditions aligns with findings that reducing spatial information impairs spatial navigation (Bates & Wolbers, 2014; Nardini et al., 2008; McCracken et al., 2025; Shayman et al., 2024). Yet, despite this increased navigational difficulty, gait and balance remained unchanged, indicating that the combined attentional demand of navigation and locomotion may not have exceeded capacity. To evaluate this explanation more directly, future work should increase the difficulty of locomotor control using perturbations that challenge vestibular and proprioceptive processing, such as uneven or compliant surfaces (Frost et al., 2015), tendon vibration (Hwang et al., 2014; Radhakrishnan et al., 2011), or galvanic vestibular stimulation (Allred et al., 2024; Gallagher et al., 2023). Similar to the effects observed with more complex cognitive tasks, increasing locomotor difficulty in dual-task paradigms reliably produces larger changes in gait, including reduced stride length, gait speed, and smoothness (Grosboillot et al., 2023; Hennah et al., 2021; Suri et al., 2023). It is therefore plausible that gait and balance would become more susceptible to interference from navigation in challenging environmental contexts, such as uneven terrain or low-light conditions. However, the present results indicate that during typical walking on stable surfaces, increased navigational difficulty does not interfere with gait or balance.

Given that walking and navigation are integrated processes, it is also possible that the standard “attentional capacity” model of dual-task interference does not capture their interaction. This model assumes that locomotion and navigation compete to draw from a shared, limited pool of resources, such that increased demand in one domain should impair performance in the other. However, body-based translational and rotational signals derived from walking are necessary for path integration (Cullen, 2019; Lappe et al., 1999; Loomis et al., 1993), meaning it is ecological for movement to facilitate effective navigation. Findings from Camp et al. (2023) highlight why distinguishing between competition and support is important. In the Walking Corsi Test (WCT), gait metrics changed as the number of blocks in the sequence increased, but this task requires encoding of a visuospatial sequence observed through another person’s movements while maintaining it in working memory – processes that are largely independent of self-motion cues. As the number of items increased, attentional demands for visuospatial working memory increase accordingly (Pratt et al., 2011). Indeed, the addition of a visuospatial working memory task (visual object tracking) leads to impairments in gait and balance in older adults (Chu et al., 2022). In contrast, the tracing phase of our VR homing task required participants to develop an allocentric representation of their own position relative to the environment while walking, integrating both visual landmarks and body-based self-motion cues. Because locomotion provides the proprioceptive, vestibular, and motor-prediction signals that underpin spatial updating, walking may facilitate navigation in this context rather than compete with it. To test whether locomotion synergistically supports navigation, future work should reduce the reliability of self-motion cues using vestibular, proprioceptive, or surface perturbations and assess whether navigation accuracy in a standard task (i.e., the Full Cues condition) disproportionately deteriorates, which would indicate that walking-generated signals actively contribute to navigation rather than compete with it.

While we were able to answer our research question and demonstrate that increasing the difficulty of navigation did not disrupt gait and balance, there were a few notable limitations of this study that impact our ability to interpret the results. First, we did not include strict single-task conditions as baselines for performance (e.g., walking without navigation; navigation without walking). For single-task walking, participants could walk between two points in the virtual environment to provide measures of gait and balance independent of navigation demands. For single-task navigation, participants could complete the VR homing task while seated using a handheld joystick (Barhorst-Cates et al., 2020, 2021). Thus, while our results support that changes in navigational cues does not impact locomotion, we cannot make conclusions about how the act of navigation impacts locomotion, or conversely, how locomotion influences navigation performance. Second, the use of a sample of young healthy adults limits application to older or clinical populations. Compared to healthy controls, older adults (Beurskens & Bock, 2012), and individuals with Parkinsons disease (Salazar et al., 2018), mild cognitive impairment (Bishnoi & Hernandez, 2021; Galrinho et al., 2025), stroke (Deblock-Bellamy et al., 2020), and mild traumatic brain injury (Fino et al., 2016, 2018; Howell et al., 2013) typically display greater dual-task interference, so it is possible navigation may interfere with gait and balance for these populations. Third, the use of integrated VR trackers is novel in biomechanics research, and while they offer accessibility and immediate spatial referencing within the virtual space, they show poorer accuracy than traditional marker-based systems such as Vicon (Merker et al., 2023). While sufficient for spatiotemporal gait metrics, such as step width, step length, and stride speed, they may be insufficiently accurate for high precision biomechanical analyses. This was not an aim of the present study; however, use of more precise marker-based motion tracking with a greater emphasis on joint kinetics and kinematics may reveal more subtle changes in locomotor control in response to increasingly difficult navigation.

## CONCLUSION

Increasing the difficulty of navigation by removing spatially informative sensory cues from the VE did not interfere with gait and balance in our VR ambulatory homing task. However, these manipulations did lead to a reliable decline in navigation performance. Therefore, in real-world navigation with changing sensory feedback, we do not believe the reliability of navigational cues directly interfere with gait and balance. We proposed various explanations for the observed results, such as a posture-first strategy, a combined attentional demand that did not exceed capacity, or a potential synergy between locomotion and navigation in real-world navigation contexts. However, the current experiment was not designed to answer these questions, so further investigation is necessary to elucidate the relationship between these interconnected motor and cognitive processes.

## ACKNOWLEDGEMENTS

The authors would like to thank Stevie Ayers-Harris, Gabriel Holm, Paula Kramer, Harry Reid, Jarem Eastman, Kento Nohara, and Alejandro Martinez for assisting in data collection.

